# Depression-like state induced by low-frequency repetitive transcranial magnetic stimulation to ventral medial frontal cortex in monkeys

**DOI:** 10.1101/2021.05.21.445094

**Authors:** Shinya Nakamura, Yodai Kishimoto, Masaki Sekino, Motoaki Nakamura, Ken-Ichiro Tsutsui

## Abstract

The medial frontal cortex (MFC), especially its ventral part, has long been of great interest with respect to the pathology of mood disorders. A number of human brain imaging studies have demonstrated the abnormalities of this brain region in patients with mood disorders, however, whether it is critically involved in the pathogenesis of such disorders remains to be fully elucidated. In this study, we conducted a causal study to investigate how the suppression of neural activity in the ventral region of the MFC (vMFC) affects the behavioral and physiological states of monkeys by using repetitive transcranial magnetic stimulation (rTMS). By using low-frequency rTMS (LF-rTMS) as an inhibitory intervention, we found that LF-rTMS targeting the vMFC induced a depression-like state in monkeys, which was characterized by a reduced spontaneous behavioral activity, increased plasma cortisol level, impaired sociability, and decreased motivation level. On the other hand, no such significant changes in behavioral and physiological states were observed when targeting the other MFC regions, dorsal or posterior. We further found that the administration of an antidepressant agent, ketamine, ameliorated the abnormal behavioral and physiological states induced by the LF-rTMS intervention. These findings indicate the causal involvement of the vMFC in the regulation of mood and affect and the validity of the LF-rTMS-induced dysfunction of the vMFC as a nonhuman primate model of the depression-like state.

## Introduction

Mood disorders are common mental illnesses characterized by sustained shifts of the vital feeling that reflects a physical aspect of affections rather than a psychological one. Nowadays, major depressive disorder has been the leading cause of disability and also a disease burden globally (1). The pathology of mood disorders and its neural background have long been studied with regards to abnormalities and variations of monoamine functions, gene expressions, and more recently, immunity and inflammation (2–4). Besides, it is quite important to understand them from the viewpoint of systems neuroscience, i.e., the malfunction of which brain region or which neural circuit is critically related to depression. In general, such approach is highly important in understanding and developing better therapeutics for mental and neurological disorders. A well-known successful example of a systems neuroscience approach would be electrophysiological studies of the basal ganglia circuit using nonhuman primates, leading to the understanding of its function and applications to deep brain stimulation (DBS) therapy in Parkinson disease (5). Some structural and functional neuroimaging studies of patients with depression showed decreases in cortical volume and abnormal activities in the medical frontal cortex (MFC), centering in the ventral anterior cingulate cortex (ACC) anterior and ventral to the genu of the corpus callosum, often referred to as the “pregenual” ACC (pgACC) and “subgenual” ACC (sgACC) (6–8). Besides these correlative lines of evidence, it is important to examine the causal involvement of the abnormalities of this area in mood-related disorders. The purpose of this study is to examine the causal relationship between the malfunction of the ventral region of the MFC (vMFC) and the expression of depression by local neural inactivation using monkeys as subjects. As for the means of local neural inactivation, we used repetitive transcranial magnetic stimulation (rTMS) (9), because it is non-invasive and yields highly reliable data from a small number of subjects through repeated measures.

## Results

In this study, we used low-frequency rTMS (LF-rTMS) as a means of local neural inactivation for two Japanese monkeys. The experimental design consisted of one experimental stimulation condition (“Exp”), in which we targeted the vMFC, and three control conditions, including two control stimulation conditions (“Con-1” and “Con-2”) and a sham stimulation condition (“Sham”). The extent of the stimulated brain areas under Exp as well as Con-1 and Con-2 was simulated using the electromagnetic model. For these four conditions, we monitored the spontaneous behavioral activity, food and water intakes, and blood plasma cortisol level in the monkeys in their home cages, and assessed their sociability and motivation. We further tested the therapeutic effects of ketamine administration on behavioral and physiological changes under Exp.

### Simulation of TMS-induced eddy current density in the monkey brain

To assess the extent of the brain areas stimulated under different stimulation conditions, we simulated the TMS-induced eddy current density in the brain model of the Japanese monkey by the scalar potential finite difference (SPFD) method (10, 11). As shown in Fig. 1C, besides the TMS induced the eddy currents in somewhat broad areas of the brain, ranging from the frontal to parietal lobes, we found different patterns of the distribution of the eddy current density among the three stimulation conditions used in this study. Under Exp, the eddy currents were induced in broad areas of the MFC and reached the vMFC (Fig. 1C). Under the control stimulation conditions, the induced eddy currents were almost restricted in the dorsal part of the MFC (Fig. 1C, Con-1), or the eddy currents were induced broadly in the dorsomedial part of the frontal and parietal cortices (Fig. 1C, Con-2), but did not reach sufficiently to the vMFC. Thus, although the distribution patterns of the eddy currents induced under these stimulation conditions widely overlapped on the mediodorsal part of the neocortex, only under Exp did the eddy currents reach the vMFC including the pgACC and sgACC.

**Figure 1.**
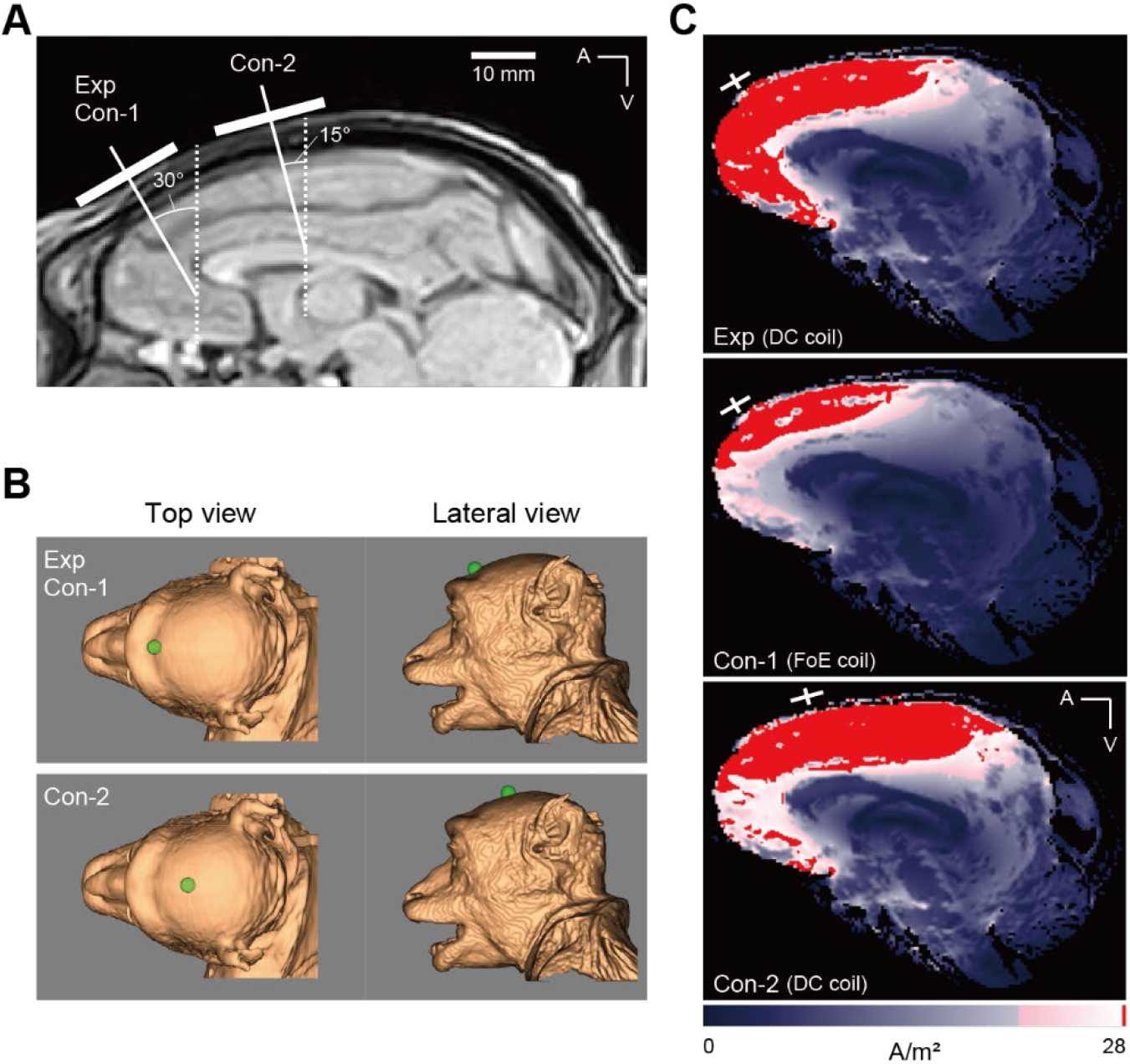
Positioning of TMS coils and simulation of induced currents under three TMS conditions (Exp, Con-1, and Con-2). (A) Sagittal MRI image of the monkey brain. White bars on the scalp represent the location and orientation of the TMS coil. (B) Images of the monkey’s head reconstructed from MR images using the TMS neuronavigation system. Green spheres on the scalp indicate the location of the center of the TMS coil. (C) Simulation of TMS-induced eddy current density in the monkey brain. White crosses indicate the location and orientation of the TMS coil. The TMS coil used for each stimulation condition is indicated in parentheses. DC; double cone, FoE: figure of eight, A: anterior, V: ventral.

### Spontaneous behavioral activity in the home cage

To investigate how the LF-rTMS intervention could affect the behavioral activity of the monkeys, we first videotaped their behavior in their home cages and evaluated the difference in their behavioral activity before and after the LF-rTMS by visual inspection. Normally (i.e., on the baseline days), the monkeys generally sat on the floor of the cage calmly and groomed themselves, sometimes looked around the surrounding, and occasionally showed whole-body movements such as walking around the floor and hanging on the cage. After the experimental stimulation, we observed a striking reduction in the level of their activeness in their behavior in the home cages for a whole day. We scaled the activeness of monkeys into five levels according to characteristic behavioral patterns from “lying” to “whole-body movements” (Figs. S1 and S2, see Materials and Methods for details). We found a significant change in the distribution of these behavioral patterns compared with that before the experimental stimulation (*P* = 6.81 × 10^−16^ and 0.00133 for monkeys A and M, respectively, Kolmogorov–Smirnov test) (Fig. 2A). Individually, we found significant decreases in the durations of whole-body movements and “sitting with hand and neck movements” (*P* = 5.98 × 10^−5^ and 5.21 × 10^−5^) and significant increases in the durations of “sitting still” and lying in monkey A (*P* = 7.70 × 10^−5^ and 6.92 × 10^−5^, respectively; paired t-test with Benjamini–Hochberg correction). In monkey M, we found significant or marginal decreases in the durations of whole-body movements, sitting with hand and neck movements, and “sitting with neck movements” (*P* = 0.0157, 0.0188, and 0.0878, respectively) and significant increases in the durations of sitting still and lying (*P* = 0.0195 and 0.0466, respectively paired t-test with Benjamini–Hochberg correction). These results indicate that their activeness shifted to low levels by the experimental stimulation. To automate the measurement of behavioral activity, we used a three-axis accelerometer attached on the neck collar of each monkey. As shown in Fig. 2B, both monkeys showed increased behavioral activity in the evening (between 16:00 to 20:00) in the normal state, whereas such an increase in behavioral activity was indeed attenuated by the experimental stimulation. We observed no such obvious changes in behavioral activity after the stimulation under the other conditions (Fig. S3). Fig. 3A shows the statistical evaluation of changes in the spontaneous behavioral activity during the daytime measured by the accelerometer. We found a significant decrease in behavioral activity on the day of the experimental stimulation, but not under the other conditions (*P* = 0.0156 and 0.0312 for Exp in monkeys A and M, respectively; *P* > 0.2 for other conditions; Wilcoxon signed-rank test with Bonferroni correction; Fig. 3A, left). This decreased behavioral activity returned to the baseline levels on the next day in monkey A, whereas it tended to continue until the next day with marginal significance in monkey M (*P* = 0.0936, Wilcoxon signed-rank test with Bonferroni correction; Fig. 3A, right).

**Figure 2.**
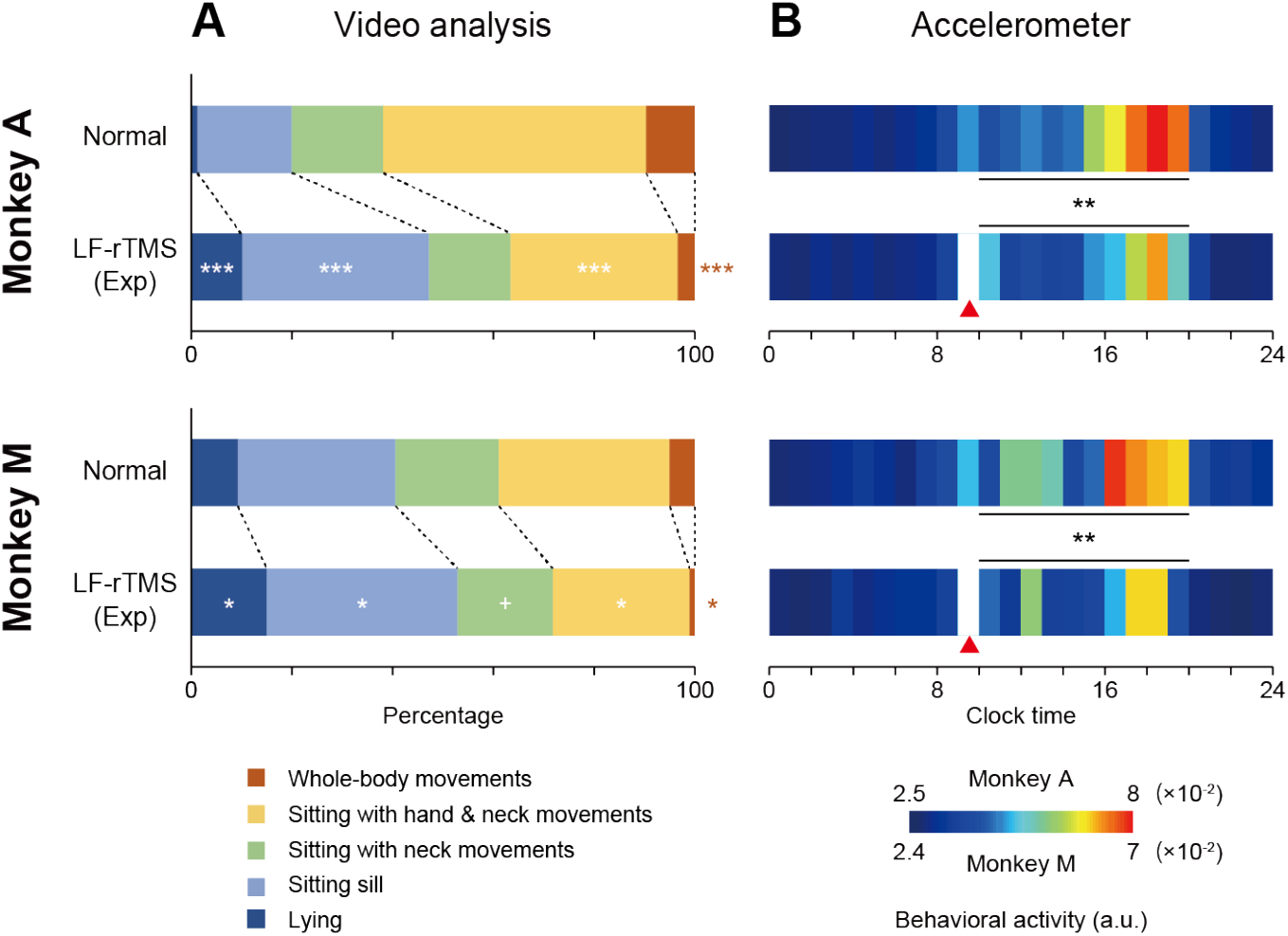
Spontaneous behavioral activity of the monkey in normal and LF-rTMS-induced depressive states. (A) Average ratios of appearance of five behavioral patterns during the daytime measured by video monitoring. Results of eight days were averaged in both monkeys. ^+^, *, \*\*\**P* < 0.1, 0.05, 0.001, paired t-test with Benjamini–Hochberg correction for multiple comparisons. (B) Time course of the spontaneous behavioral activity in the home-cage measured by the accelerometer. Red arrowheads indicate the timing of LF-rTMS intervention. Results of nine and eight days were averaged in monkeys A and M, respectively. \*\**P* < 0.01, Wilcoxon signed-rank test.

**Figure 3.**
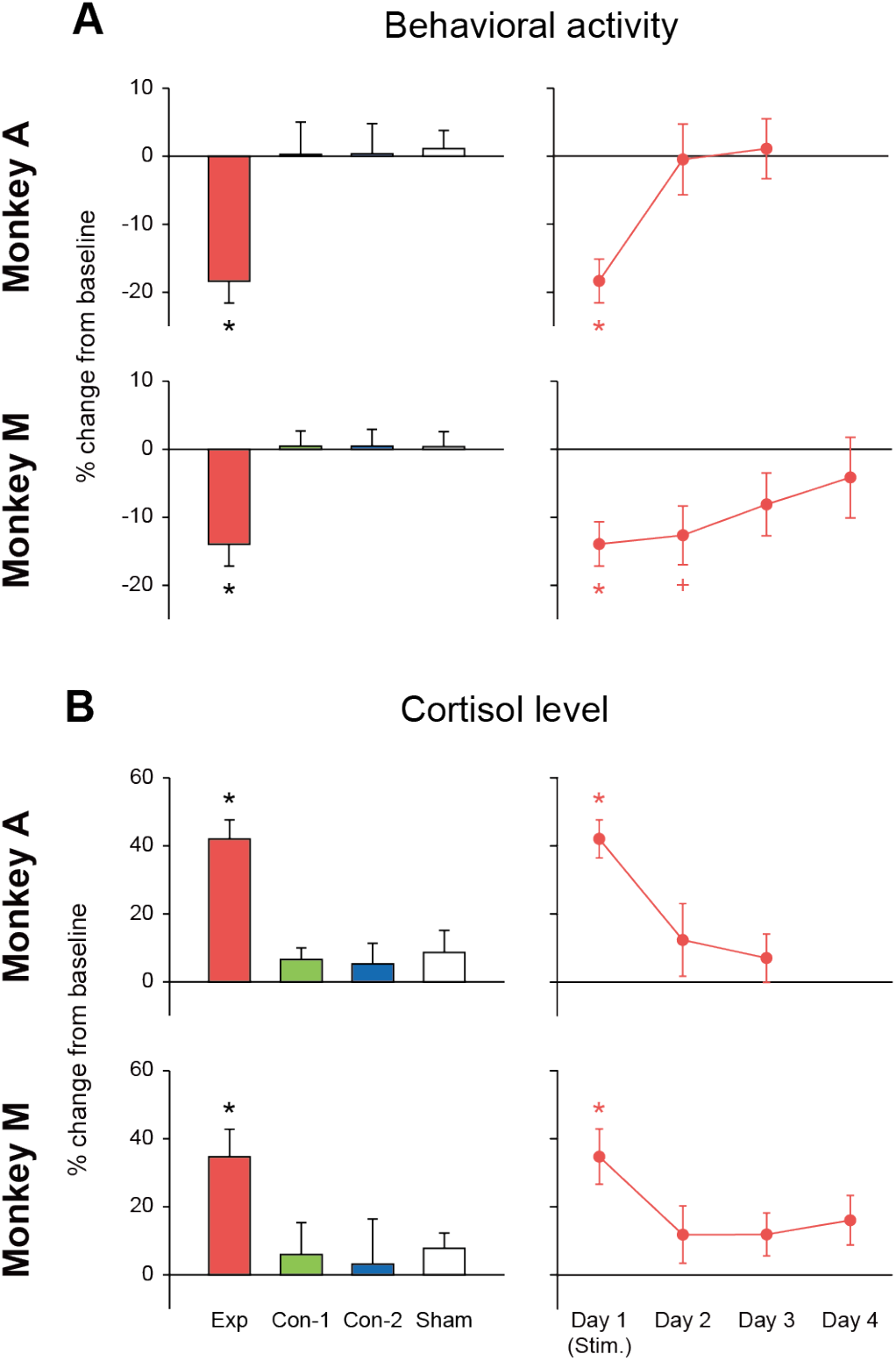
Effects of LF-rTMS intervention on the behavioral and physiological states of the monkey. The percent changes in the spontaneous behavioral activity in the home-cage (A) and evening plasma cortisol level (B) compared with the baseline levels are shown. Bar graphs show the results of the day we performed the stimulation. Line graphs show the time course of the behavioral and physiological changes under Exp. The data are shown as mean ± SEM. N = 9, 7, 7, and 7 (monkey A) and 8, 6, 6, and 6 (monkey M) for Exp, Con-1, Con-2, and Sham, respectively, in both A and B. ^+^, \**P* < 0.1, 0.05, Wilcoxon signed-rank test with Bonferroni correction for multiple comparisons.

### Food and water intakes

We examined changes in food (the number of pellets (approximately 2.5 g/pellet)) and water (mL) intakes (Fig. S4). In monkey A, we found a significant decrease in food intake and a slight increase in water intake under Exp (*P* = 0.0289 and 0.0731), but not under the other stimulation conditions (*P* > 0.2 for all, t-test with Bonferroni correction) when compared with the baseline levels. In monkey M, we found no significant changes in food and water intakes under any stimulation conditions (*P* > 0.2 for all, t-test with Bonferroni correction).

### Blood cortisol level

We examined the changes in evening blood plasma cortisol level under each stimulation condition (Fig. 3B). The cortisol level under Exp significantly increased with reference to the baseline measured in the evening the day before, whereas that under other stimulation conditions did not (*P* = 0.0117 and 0.0312 for Exp in monkeys A and M, respectively, *P* > 0.2 for other conditions, Wilcoxon signed-rank test with Bonferroni correction; Fig. 3B, left). The cortisol level on the next day returned to a level where there was no significant difference from the baseline level (*P* > 0.2 for both monkeys, Wilcoxon signed-rank test with Bonferroni correction; Fig. 3B, right).

### Sociability

Normally, the monkeys were sociable when an experimenter entered the animal room and approached their home cages. Both monkeys immediately approached the front side of the cages and sat close to the experimenter with their bodies and/or faces oriented towards the experimenter, but looking away and avoiding too much eye contact. Generally, monkeys are believed to avoid making too much eye contact as it can be a sign of aggression (12, 13). Besides, monkey A occasionally looked at the experimenter, which could be an affiliative sign in female monkeys (12), whereas monkey M occasionally showed rapid glances toward the experimenter, which could be a submissive sign in male monkeys (13). In contrast, after the experimental stimulation, both monkeys were less sociable than in their normal states. Although monkey A immediately approached the front side of the cage when it noticed the approach of the experimenter, it went back to the back side of the cage after a while. Monkey M did not immediately come to the front side of the cage and only after some time did it slowly move towards the front side. When they sat on the front side of the cage close to the experimenter, they oriented their bodies and/or faces away from the experimenter, seemingly avoiding any eye contact. Indeed, these behavioral changes made an overall impression to the experimenter that the monkeys were “depressed” and “withdrawn”. To quantitatively evaluate such behavioral changes, we performed a sociability test, which quantified the monkeys’ behavior when approached by the experimenter (Figs. 4A and 4B). By using Dunnett’s multiple comparison test, we found a significant increase in the latency under Exp in monkey M (*P* = 6.96 × 10^−5^) but not in monkey A (*P* = 0.195, Fig. 4C). The percent time in the interaction zone significantly decreased under Exp in both monkeys (*P* = 0.00786 and 1.24 × 10^−4^ for monkeys A and M, respectively; Fig. 4D). The percent time the monkeys oriented their faces toward the experimenter significantly decreased under Exp in monkey A (*P* = 0.00177), but not in monkey M (*P* = 0.946, Fig. 4E). The percent time the monkeys oriented their bodies toward the experimenter significantly decreased in monkey A (*P* = 0.00360) and marginally decreased in monkey M (*P* = 0.0501) under Exp (Fig. 4F). The percent time the monkeys showed straightly looking-down posture significantly increased under Exp in both monkeys (*P* = 1.51 × 10^−5^ and 0.00247 for monkeys A and M, respectively; Fig. 4G). Although the induced behavioral changes under Exp were slightly different between the two monkeys, it may be reasonable to interpret that the level of sociability was significantly reduced in both monkeys after the experimental stimulation.

**Figure 4.**
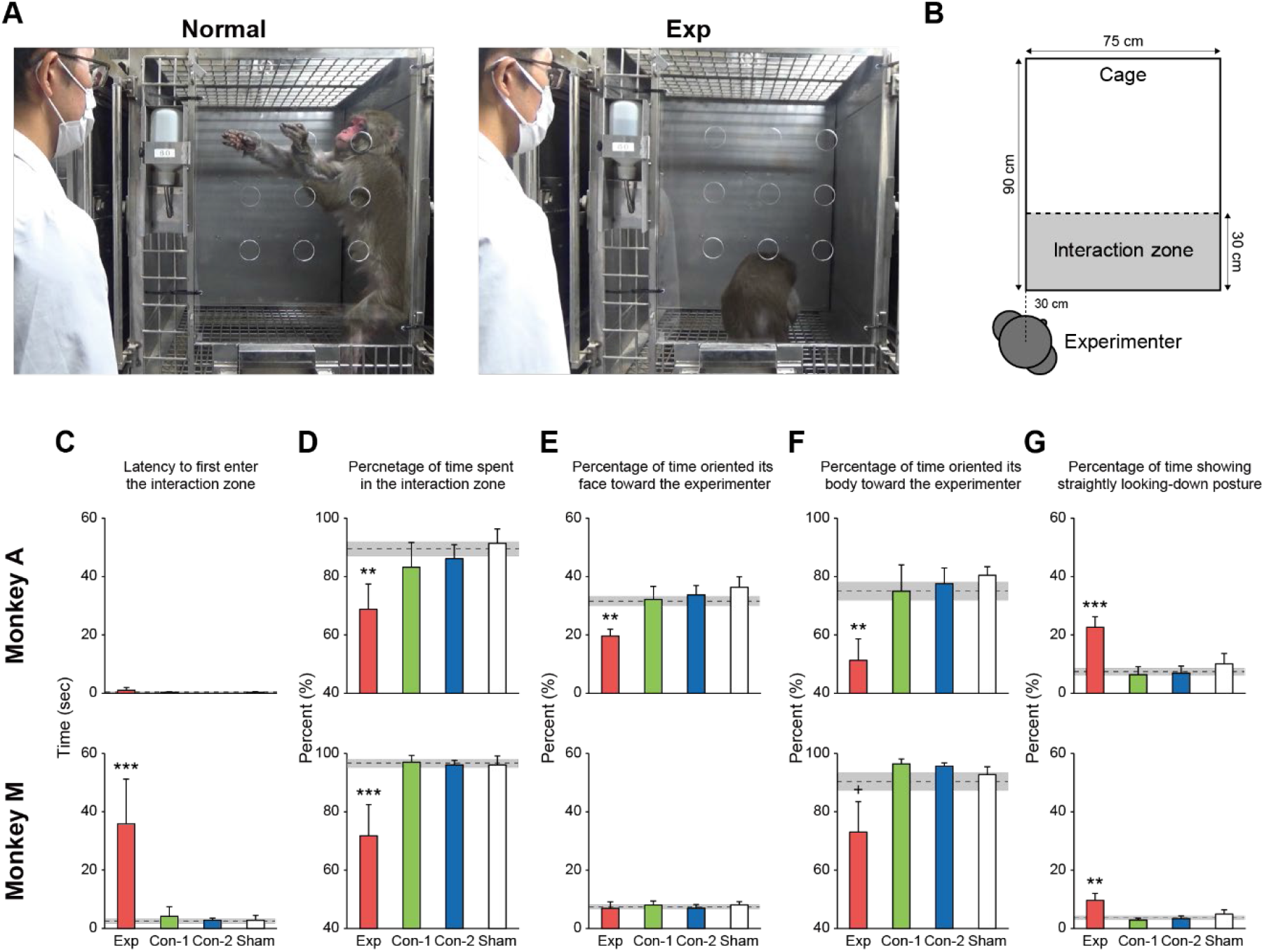
Effects of the LF-rTMS intervention on monkey sociability. (A) Pictures of the monkeys during the social interaction test. The monkeys were normally sociable and approached the front side of the cage close to the experimenter (left), whereas under Exp, the monkeys were less sociable and withdrawn (right). (B) Illustration showing the locations of the experimenter and the interaction zone in the sociability test. (C) Latency to first enter the interaction zone. (D) Percentages of the time spent within the interaction zone. (E) Percentages of the time the monkey oriented its face toward the experimenter. (F) Percentages of the time the monkey oriented its body toward the experimenter. (G) Percentages of the time showing straightly looking-down posture. Gray dotted lines indicate baseline levels. The data are shown as mean ± SEM (error bars and shaded areas). N = 9, 7, 7, and 7 (monkey A) and 8, 6, 6, and 6 (monkey M) for Exp, Con-1, Con-2, and Sham, respectively. ^+^, **, \*\*\**P* < 0.1, 0.01, 0.001, Dunnett’s multiple comparison test.

### Motivation

We examined the change in the motivational level of the monkeys following each stimulation by using a modified version of the Brinkman board test (14–16), in which we introduced two difficulty levels of the test. We quantified the number of sessions the monkey performed until it spontaneously stopped performing and the average time that the monkey required to complete a single session as indices of the motivation and motor functionality, respectively (Fig. 5, see Materials and Methods for details). Two-way analysis of variance (ANOVA) revealed a significant main effect of the difficulty level on the number of trials under all stimulation conditions (*F*_(1,20)_ = 64.14, *P* = 1.14 × 10^−7^ (Exp); *F*_(1,16)_ = 10.58, *P* = 0.00498 (Con-1); *F*_(1,16)_ = 29.68, *P* = 5.36 × 10^−5^ (Con-2); *F*_(1,16)_ = 26.65, *P* = 9.44 × 10^−5^ (Sham) for monkey A; *F*_(1,16)_ =127.19, *P* = 5.02 × 10^−9^ (Exp); *F*_(1,16)_ = 9.34, *P* = 0.00752 (Con-1); *F*_(1,16)_ = 18.33, *P* = 5.72 × 10^−4^ (Con-2); *F*_(1,16)_ = 16.66, *P* = 8.68 × 10^−4^ (Sham) for monkey M). We further found that the number of trials significantly decreased after the experimental stimulation when the monkeys performed the difficult test (*F*_(1,20)_ = 11.78 and *P* = 0.00263 (interaction), *P* = 0.00104 (post hoc) for monkey A; *F*_(1,16)_ =9.19 and *P* = 0.00793 (interaction), *P* = 0.00237 (post hoc) for monkey M). Regarding the average time, we only found significant main effects of difficulty level under all stimulation conditions (*F*_(1,20)_ = 65.66, *P* = 1.38 × 10^−7^ (Exp); *F*_(1,16)_ = 52.96, *P* = 1.85 × 10^−6^ (Con-1); *F*_(1,16)_ = 32.64, *P* = 3.20 × 10^−5^ (Con-2); *F*_(1,16)_ = 39.46, *P* = 1.09 × 10^−5^ (Sham) for monkey A; *F*_(1,16)_ = 151.01, *P* = 1.45 × 10^−9^ (Exp); *F*_(1,16)_ = 17.05, *P* = 7.86 × 10^−4^ (Con-1); *F*_(1,16)_ = 48.11, *P* = 3.35 × 10^−6^ (Con-2); *F*_(1,16)_ = 87.24, *P* = 7.03 × 10^−8^ (Sham) for monkey M). These results indicate that the decrease in the number of sessions performed in the difficult test induced by the experimental stimulation may not be attributed to the disturbance of motor function, and should be attributed to the decrease in motivational level.

**Figure 5.**
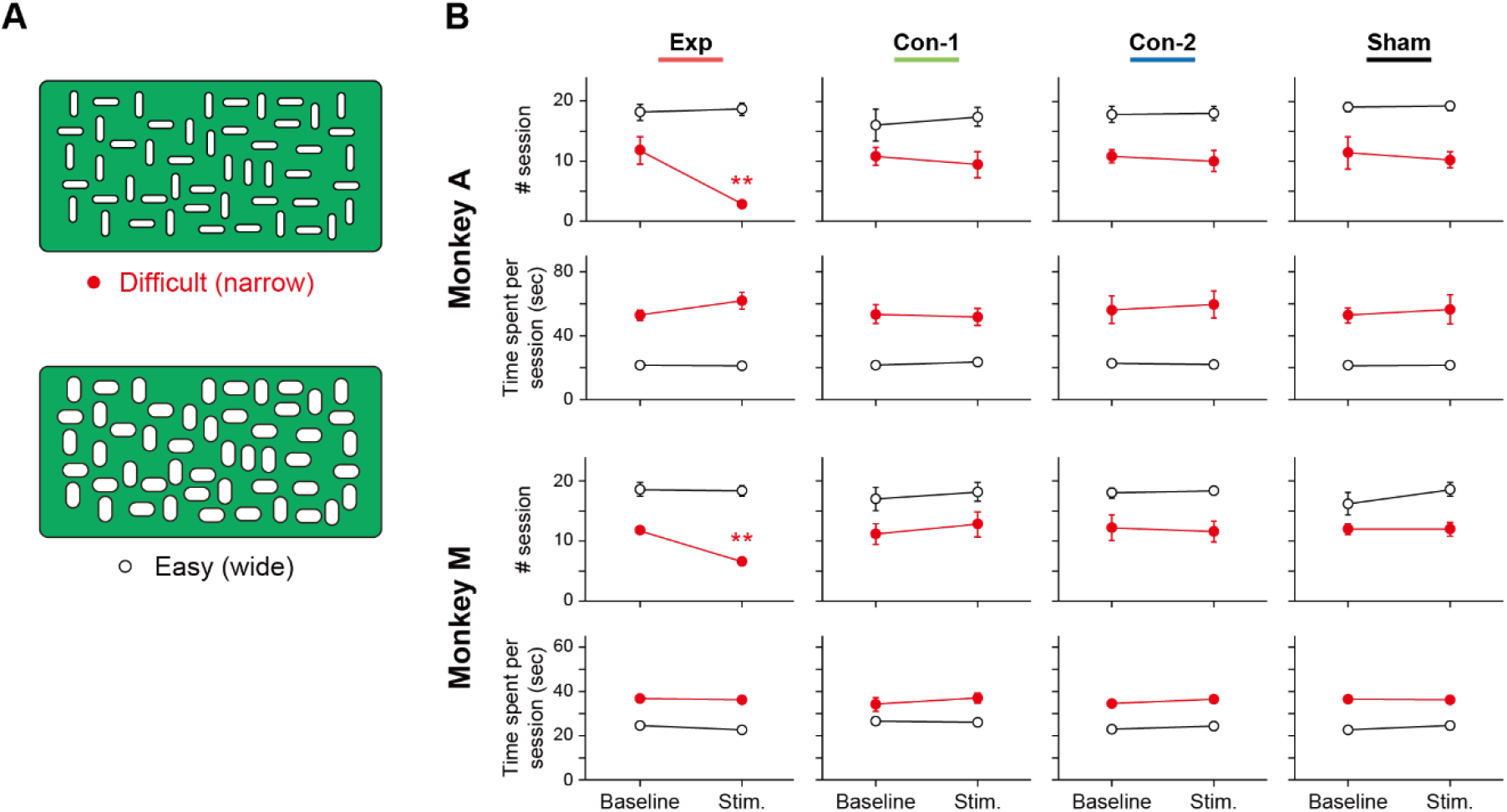
Effects of LF-rTMS intervention on the monkeys’ motivational level. (A) Illustrations of modified Brinkman boards with narrow (difficult, top) and wide (easy, bottom) wells. (B) Changes in the performance of the motivational test with the Brinkman board. The number of sessions performed until the monkey spontaneously stopped performing (upper graphs) and the average time required to finish a single session (lower graphs) are shown for each monkey. The data are shown as mean ± SEM. N = 6, 5, 5, and 5 (monkey A) and 5, 5, 5, and 5 (monkey M) for Exp, Con-1, Con-2, and Sham, respectively, in both difficult and easy conditions. \*\**P* < 0.01, two-way ANOVA with post-hoc Bonferroni analysis.

### Effects of ketamine administration

The above results led to the question on whether the behavioral and physiological changes induced by LF-rTMS under Exp could be normalized by antidepressant treatments. To answer this question, we chose ketamine as an antidepressant because recent preclinical studies have demonstrated that a single intravenous administration of sub-anesthetic doses of ketamine has rapid and robust antidepressant effects in patients with MDD manifesting within a few hours (17–19). We examined the spontaneous behavioral activity and evening plasma cortisol level in the monkeys intravenously administered the low-dose ketamine (0.5–1.0 mg/kg, see Materials and Methods for details) after the experimental stimulation (Exp+Keta) or without the LF-rTMS intervention (Keta). First, the administration of ketamine did not induce any obvious behavioral or physiological acute reactions, such as nystagmus or salivation. Besides, as shown in Fig. 6, we found that the reduced spontaneous behavioral activity and increased evening plasma cortisol level induced by the experimental stimulation were significantly improved by the ketamine administration, and that the behavioral activity and cortisol level under the Exp+Keta and Keta conditions were not significantly different from those under the Sham condition (*P* = 0.0213 and 0.0204 (behavioral activity), and *P* = 0.0191 and 0.0387 (cortisol) for Exp vs. Exp+Keta in monkeys A and M, respectively; *P* > 0.2 for Exp+Keta vs. Sham and Keta vs. Sham in both monkeys; t-test with Bonferroni correction). Considering that the intravenous administration of low-dose ketamine itself did not significantly affect the behavioral and physiological states of the monkeys, these results indicate that the abnormal state induced by the experimental stimulation was ameliorated by the antidepressant effects of the ketamine.

**Figure 6.**
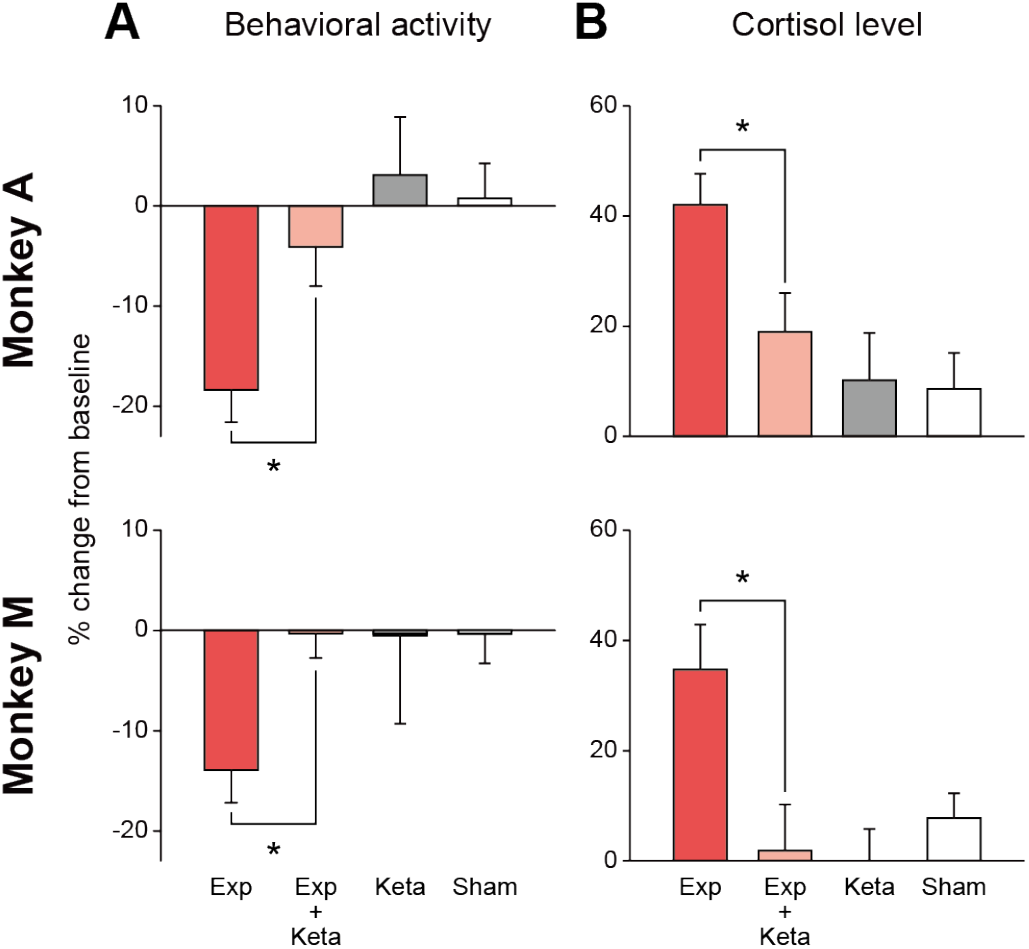
Effects of intravenous administration of ketamine on abnormal behavioral and physiological states induced by LF-rTMS intervention under Exp. (A) Percent changes in spontaneous behavioral activity in the home cage compared with the baseline level (B) Percent changes in evening plasma cortisol level compared with the baseline level. The data are shown as mean ± SEM. Exp+Keta: ketamine administration following LF-rTMS intervention under Exp. Keta: ketamine administration without stimulation. N = 9, 6, 6, and 7 (monkey A), and 8, 6, 7, and 6 (monkey M) for Exp, Exp+Keta, Keta, and Sham, respectively. \**P* < 0.05, t-test with Bonferroni correction for multiple comparisons.

## Discussion

The present study demonstrated that LF-rTMS targeting the vMFC including the pgACC and sgACC induced a depression-like state in the monkeys, which was characterized by significant decreases in the spontaneous behavioral activity, sociability, and motivational level, and a significant increase in blood plasma cortisol level. These symptoms are consistent with those observed in patients with MDD, such as fatigue, impairment of social functioning, and loss of interest or pleasure, as listed in the fifth edition of the Diagnostic and Statistical Manual of Mental Disorders (DSM-5). Hyperactivity of the hypothalamic–pituitary–adrenal axis and hypercortisolism have also been reported in patients with MDD (20, 21). Furthermore, the administration of an antidepressant agent, ketamine, normalized the abnormalities of the spontaneous behavioral activity and the blood plasma cortisol level induced by LF-rTMS. No such symptoms were observed under the other stimulation conditions in which the eddy currents induced by TMS pulses did not sufficiently reach the vMFC. These findings indicate that the vMFC is critically involved in the regulation of mood and affect, and that dysfunction of the vMFC can cause depression.

To the extent of our knowledge, the present study demonstrated the first-ever monkey depressive-state model induced by direct neural interventions. To date, there have been several studies of the depressive state in monkeys induced spontaneously (22–25) or by behavioral (26, 27) or systemic pharmacological (28, 29) interventions. In rodents, chronic mild stress, social defeat, and pharmacological models have been generally used to develop and assess new therapeutic agents and to examine the neurobiological mechanisms of depression (30–33). However, there has been a long debate on how much the rodent depression model can be comparable to human depression, as the drugs effective for curing the rodent depression model are not necessarily effective for curing human depression (34). We consider that the monkey depression model has great advantages over existing rodent models, especially for the systems neuroscience approach, because of the high homology of brain structures, such as the ACC and the surrounding frontal cortices, between humans and monkeys.

The vMFC has long been implicated in the regulation of mood and affect (35–37). In particular, physiological and morphological abnormalities of the pgACC and sgACC in patients with MDD have been consistently observed in human imaging studies (6–8). Human clinical studies have also shown that high-frequency DBS of the sgACC can improve depressive symptoms in patients with treatment-resistant depression (35–37, but see 38). Some nonhuman primate studies have shown the involvement of these MFC areas in anxiety and anhedonia, which are widely recognized as major symptoms of MDD, by using sophisticated behavioral paradigms. In marmoset monkeys, pharmacological inactivation of area 25 (sgACC) induced the attenuation of anticipatory negative emotional responses and the enhancement of fear extinction, whereas the inactivation of area 32 (pgACC) induced the opposite effects during the fear conditioning task (42). Pharmacological overactivation of the marmoset sgACC attenuated appetitive anticipatory arousal and the reward motivation (43). On the other hand, in macaque monkeys, sgACC lesioning impaired the maintenance of autonomic arousal during anticipation of rewards during the Pavlovian conditioning task (44), and the microstimulation of the pgACC induced an increase in the frequency of avoidance decisions during the performance of the approach–avoidance task (45). In this study, however, we did not directly assess the anxiety and anhedonia symptoms using such behavioral paradigms. Nevertheless, there has been no study reported so far showing drastic changes in spontaneous and social behaviors that highly resemble the symptoms reported as naturally induced depression in monkeys (22–25).

Generally, neuromodulatory effects of a single rTMS intervention session depend on the number of pulses and the stimulation intensity, whereas the duration and size of the effects are limited (46–48). In this study, the monkeys were still willing enough to perform the motivational test on a chair even when in the depressive state induced by LF-rTMS targeting the vMFC, and they recovered from such state in a day or two, indicating that the depressive state observed in this study is moderate and acute. However, such limited effectiveness of a single rTMS intervention can be enhanced by repetition, possibly through neural plasticity mechanisms (47). Indeed, in our preliminary observation, 3-4 repetitive sessions (one session/day) of LF-rTMS targeting the vMFC induced severer and longer-lasting augmented depressive state in monkeys (Fig. S5). In such state, those monkeys spent much of their time looking down with a hunched posture, as reported in the cases of naturally-occurring depression observed in socially-housed monkeys (22–25).

As TMS/rTMS has trans-synaptic large-scale network effects on brain activity beyond the stimulated brain region (49), one interpretation of the current results is that LF-rTMS targeting the vMFC including the pgACC and sgACC modulated the activity of other brain regions functionally and/or anatomically connected to these brain regions and induced the malfunction of whole-brain networks that could be related to the regulation of mood and affect, which causes the depressive symptoms observed. Indeed, human brain imaging studies have shown a significant difference in functional connectivity in the default mode network, a resting-state functional network mediating self-referential processing (50), between patients with MDD and healthy controls (51, 52). As high-frequency rTMS to the left dorsolateral prefrontal cortex (DLPFC) has been established and widely used as an effective rTMS treatment for depression (53, 54), it is thought that the DLPFC stimulation can modulate the function of the vMFC and other limbic regions and retune the activity of prefrontal and limbic networks (53, 55). Anatomically, the pgACC and sgACC have strong fiber connections with cortical and subcortical regions, such as the orbitofrontal cortex (56), amygdala (57), nucleus accumbens (58), and hypothalamus (59), which are all considered to be involved in emotional processes (60). Considering such anatomical connections, it is conceivable that these neural circuits differentially contribute to emotional, social, and motivational behaviors, and that the malfunction of the vMFC totally disrupted the normal functioning of the brain network that consists of these neural circuits. However, it is technically difficult to stimulate and modulate the activity of individual neural circuits by rTMS. Recent development of genetic tools, such as opto- and chemogenetics, enables researchers to bidirectionally modulate the activity of genetically targeted neurons and/or brain circuits (61). By using these techniques, researchers should address in future studies how such functional and anatomical connections differentially contribute to the emotional, social, and motivational behaviors and how the malfunctions of them are related to different aspects of the symptoms of depression.

## Materials and Methods

### Subjects

Two Japanese monkeys (*Macaca fuscata*) (monkey A: female, 14–15 years of age, 8.64 ± 0.20 kg, monkey M: male, 10–11 years of age, 11.33 ± 0.53 kg, during experiment periods) served in this study. They were housed individually in appropriate cages (approximately W750 × D900 × H900 mm) in a room with a 12:12 light dark cycle (lights on at 0800). All procedures were conducted in accordance with the National Institutes of Health’s *Guide for the Care and Use of Laboratory Animals* and the Tohoku University’s *Guidelines for Animal Care and Use*. This project was approved by the Center for Laboratory Animal Research of Tohoku University.

### rTMS

LF-rTMS (1 Hz, 1200 pulses in total) was delivered to subregions of the monkey MFC with a Magstim Rapid2 stimulator with a 70-mm double cone (DC) coil and a 70-mm figure-of-eight (FoE) coil (The Magstim Company Ltd., UK). Four stimulation conditions were used in this study: (1) the DC coil was placed on the scalp with its center located on the midline 49 mm anterior to the interaural line and its surface slanted approximately 30° towards the front (Exp), (2) the FoE coil was placed on the same place and with the same orientation as Exp (Con-1), (3) the DC coil was placed on the scalp with its center located on the midline 26 mm anterior to the interaural line and its surface slanted approximately 15° towards the front (Con-2), and (4) 1200 pulses of a brief electrical current (50 V, 200 μs) were delivered to the electrode placed on the scalp at 1 Hz synchronized with a clicking sound of a TMS pulse (sham stimulation). The TMS coil was placed on the scalp so that a biphasic current was induced with an initial posteroanterior direction. The head of the monkey was firmly but noninvasively fixed during the stimulation with a head holder made from splint materials (Polyflex Ⅱ) and customized to fit the shape of the head of each monkey. The intensity of TMS was set to 150% of the resting motor threshold (rMT) of each monkey. The rMT of each monkey was defined as the machine output that produced a visible twitch in its foot in 5 out of 10 TMS pulses delivered to the primary motor cortex while the monkey calmly sat on the monkey chair. The coil position and orientation were determined using the frameless stereotaxic system (Brainsight 2, Rogue Research, Montreal, Canada). The rTMS sessions were conducted between 9:00 AM and 11:00 AM.

### Spontaneous behavioral activity in the home cages

The spontaneous behavioral activity of monkeys in their home cages was assessed by video monitoring and accelerometer measurement. For the video monitoring analysis, the behaviors of the monkeys monitored by a standard USB camera (Logicool) during daytime were categorized mainly into five patterns: whole-body movements (e.g., hanging on the cage, standing, and walking), sitting with hand and neck movements, sitting with neck movements, sitting still, and lying down on the floor of the cage. To estimate the frequency of appearances of these five behavioral patterns during daytime, we counted the number of times each behavioral pattern appeared at 480 time points (approximately every one minute) after the stimulation session. The corresponding time period was used for the analysis of baseline data.

For the accelerometer measurements, a three-axis accelerometer was attached to the loose-fitting neck collar of each monkey. Data were transmitted wirelessly to and stored in the computer (sampling rate: 10 Hz) through a home-made software developed in Visual C++ (Microsoft Visual Studio 2008). To remove the gravitational component from the raw acceleration data, we calculated the behavioral activity of the monkeys as follows:

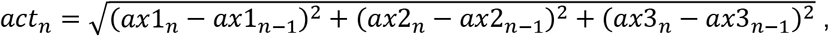

where *ax1_n_*, *ax2_n_*, and *ax3_n_* are the raw accelerometer data of three axes, *act_n_* is a calculated behavioral activity, and *n* refers to the accelerometer *n*th sample. *act_n_* is called ‘jerk’, the derivative of acceleration. The jerk can be an alternative value for estimating behavioral activity, and its use enables the easy removal of constant offsets, i.e., the gravitational component, from the raw acceleration data (62). As shown in Fig. S2, the values were high when the monkeys were showing active behaviors (whole-body movements, sitting with hand and neck movements, and sitting with neck movements), whereas those were low when the monkeys were keeping still (sitting still or lying).

### Cortisol level

Blood samples were collected from a forelimb (monkey M) or hindlimb (monkey A) in the evening (17:00–18:00 hours) without sedation to determine plasma cortisol level.

### Sociability test

To evaluate the sociability of the monkeys, we performed the sociability test, in which their behavior toward the experimenter in the home cages was examined. The behavior of the monkeys during the test was monitored with a USB camera. The size of the cage was 75 cm wide, 90 cm deep, and 90 cm high, and the experimenter (SN) stood approximately 30 cm away from the left corner of the cage for approximately 2 min (Fig. 4B). During the sociability test, the experimenter looked at the back side wall of the cage and occasionally reached out to the cage. One-third (75 × 30 × 90 cm) of the home cage area from the side near the experimenter was defined as the interaction zone (Fig. 4B). To quantify the sociability of the monkeys, we measured the latency to first enter the interaction zone, the time that the monkeys spent within the interaction zone, the time that the monkeys oriented their faces and bodies toward the experimenter, and the time that the monkeys showed straightly looking-down posture.

### Motivation

To quantified the motivational level of the monkeys, we introduced a behavioral paradigm with a modified version of the Brinkman board, which was originally developed to test manual dexterity (11–13). The board (14 × 22 cm) consisted of 25 vertically and 25 horizontally oriented wells. Each well was 15 mm long, 5 mm deep, and 4-8 mm wide (Fig. 5A). The board was set in front of the monkey sitting in a primate chair at approximately 20° tilt from the horizontal plane. The test required the monkey to grasp a small food cube 3–4 mm on each side (sweet potatoes or dried bananas) placed in each well. The food cubes were randomly placed in 12 wells per session (6 vertical and 6 horizontal wells). Two types of boards were used to change the difficulty of the test for each monkey: one had narrow wells (difficult, 4.5 mm for monkey A and 4 mm for monkey M) and the other had wide wells (easy, 6 mm for monkey A and 8 mm for monkey M). Multiple sessions were continued until the monkey spontaneously stopped performing (i.e., giving up) the session. Each session lasted up to 2 min. Each session was interrupted when one of the following cases occurred: (1) the monkey did not start the session within 30 sec, (2) the monkey stopped to grasp the food cubes for over 30 sec, or (3) 2 min had passed from the beginning of the session. In the first two cases, the monkey was considered to have given up the session, and the test was terminated. In the last case, the time the monkey spent during the session was excluded from the calculation (see below). Two parameters were quantified before and after the rTMS intervention: the number of sessions that the monkey performed before the test was terminated as the index of the motivational level, and the average time that the monkey spent to complete a single session as the index of the motor functionality.

### Ketamine administration

Human preclinical studies have shown that an intravenous administration of sub-anesthetic doses of ketamine (0.5 mg/kg) over 40 min has rapid and robust antidepressant effects on patients with MDD (17). To mimic this procedure and apply it to monkeys, ketamine hydrochloride was intravenously administered four separate times with 10-min intervals (0.1 ml/infusion, total dose of 0.5–1.0 mg/kg). The first administration began at approximately 30 min after the LF-rTMS session.

### Simulation of TMS-induced eddy current density in monkey brain

The extent of TMS-induced eddy currents in the monkey brain was assessed by the SPFD method (Fig. 1C) (10, 11). We used the monkey brain model (*the MRI standard brain of Japanese macaque monkey*, http://brainatlas.brain.riken.jp/jm/modules/xoonips/listitem.php?index_id=9) (63) segmented into three materials (white matter, gray matter, and cerebrospinal fluid) and calculated the density of eddy currents induced by TMS with the intensity of 60% of the machine power.

## Acknowledgments

This study was supported by “Integrated Research on Neuropsychiatric Disorders and Development of BMI Technologies for Clinical Application” carried out under the Strategic Research Program for Brain Sciences from the Japan Agency for Medical Research and Development, Grants-in-Aid for Scientific Research on Innovative Areas “Adaptive Circuit Shift” (26112009) and “Hyper-Adaptability” (19H05725) from the Ministry of Education, Culture, Sports, Science and Technology, Japan; and Grants-in-Aid for Scientific Research (24223004, 26560455, 17H01014, 20H00104) from the Japan Society for Promotion of Science to K-I.T.

## Notes

### Competing Interest Statement

The authors have declared no competing interest.

