## Supplemental Information for "Depression-like state induced by low-frequency repetitive transcranial magnetic stimulation to ventral medial frontal cortex in monkeys"

Ken-Ichiro Tsutsui

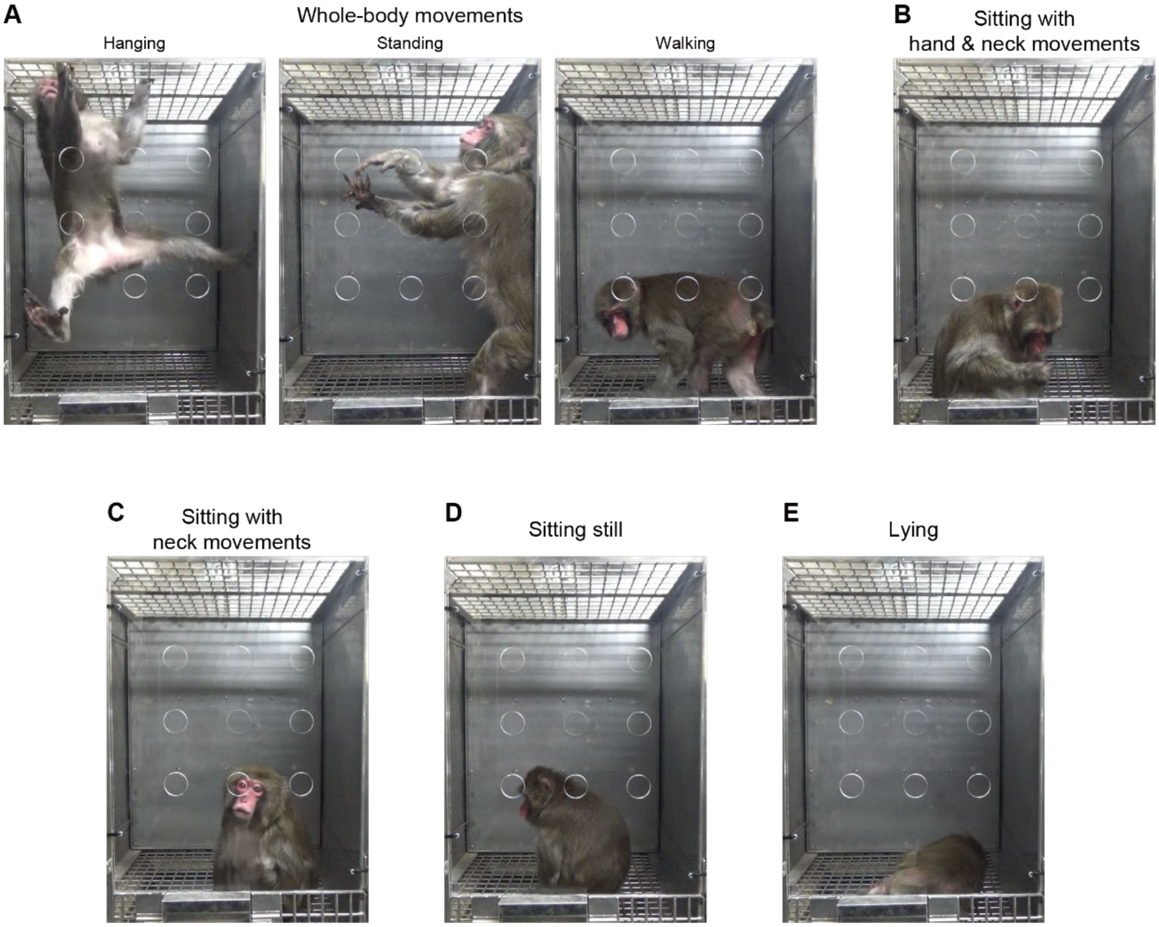


**Fig. S1** Examples of photographs of characteristic behavioral patterns that the monkeys showed in their home cages. We scaled the activeness of monkeys into five levels. (A) Whole-body movements (e.g., hanging on the cage, standing, and walking around). (B) Sitting with hand and neck movements (e.g., grooming). (C) Sitting with neck movements (e.g., looking around). (D) Sitting still (e.g., gazing somewhere). (E) Lying down on the floor.

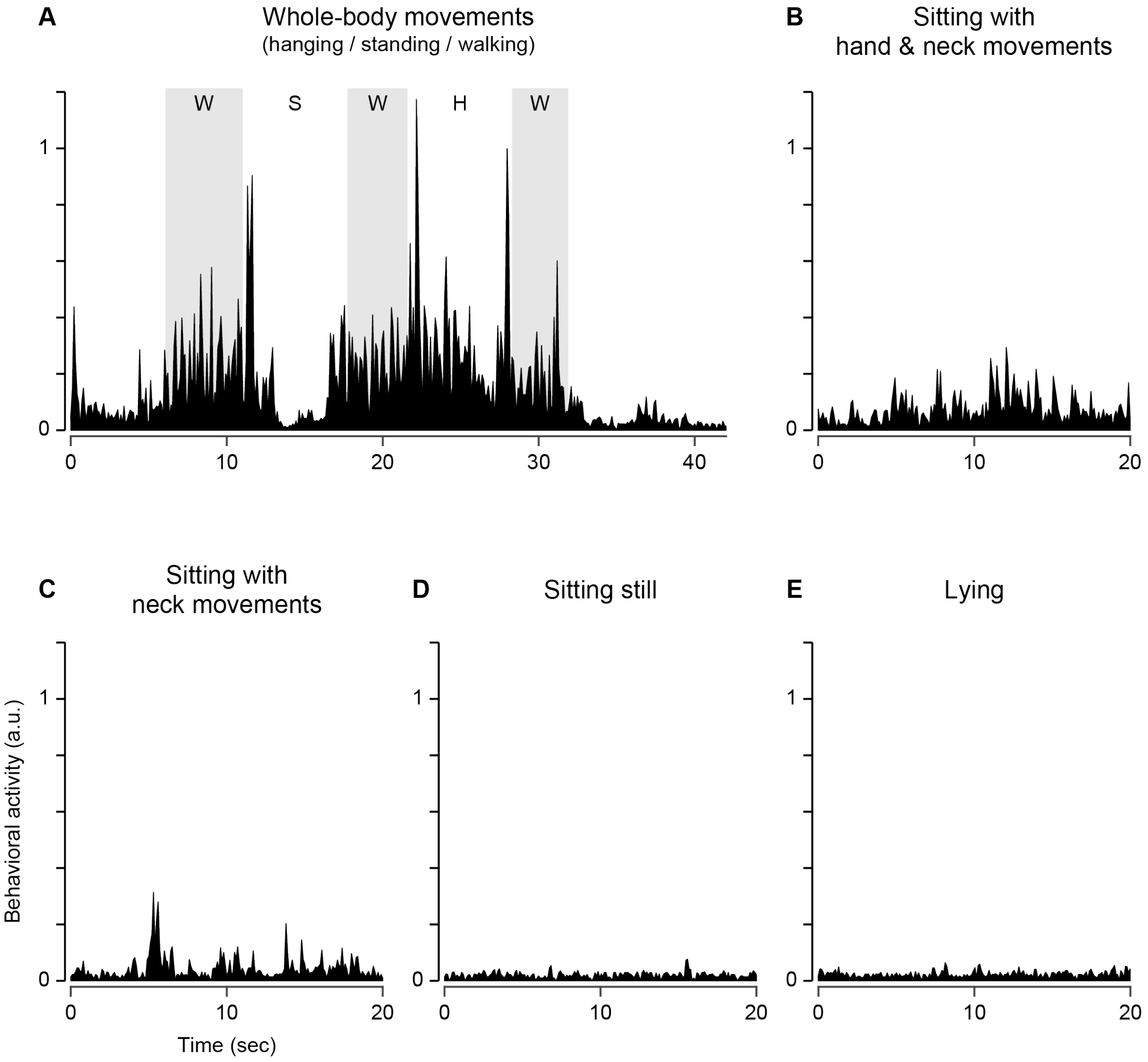


**Fig. S2** Examples of changes in behavioral activity calculated from the accelerometer values (see Materials and Methods for details) when the monkey showed different behavioral patterns. (A) Drastic increases in behavioral activity were observed when the monkey showed whole-body movements. (B and C) The behavioral activity was moderately high when the monkey was sitting with some body movements (hand and neck movements). (D and E) The behavioral activity was low when the monkey was sitting still (D) or lying on the floor (E). Note that whole-body movements, such as hanging (H), standing (S), and walking (W), usually appeared continuously. a.u., arbitrary units.

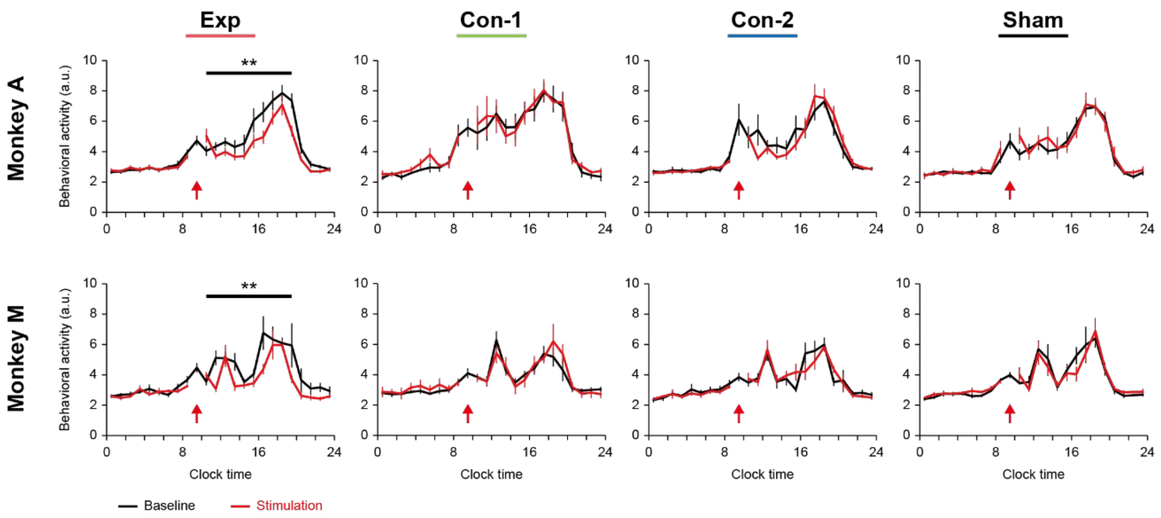


**Fig. S3** Time course of readouts of spontaneous behavioral activity from the accelerometer before and after the treatments under each condition. Red arrows indicate the timing of the stimulation. The data are shown as mean ± SEM. The behavioral activity during daytime after the stimulation (black bars) was compared between the baseline and stimulation days. N = 9, 7, 7, and 7 (monkey A), and 8, 6, 6, and 6 (monkey M) for Exp, Con-1, Con-2, and Sham, respectively. $^{**}P < 0.01$, Wilcoxon signed-rank test.

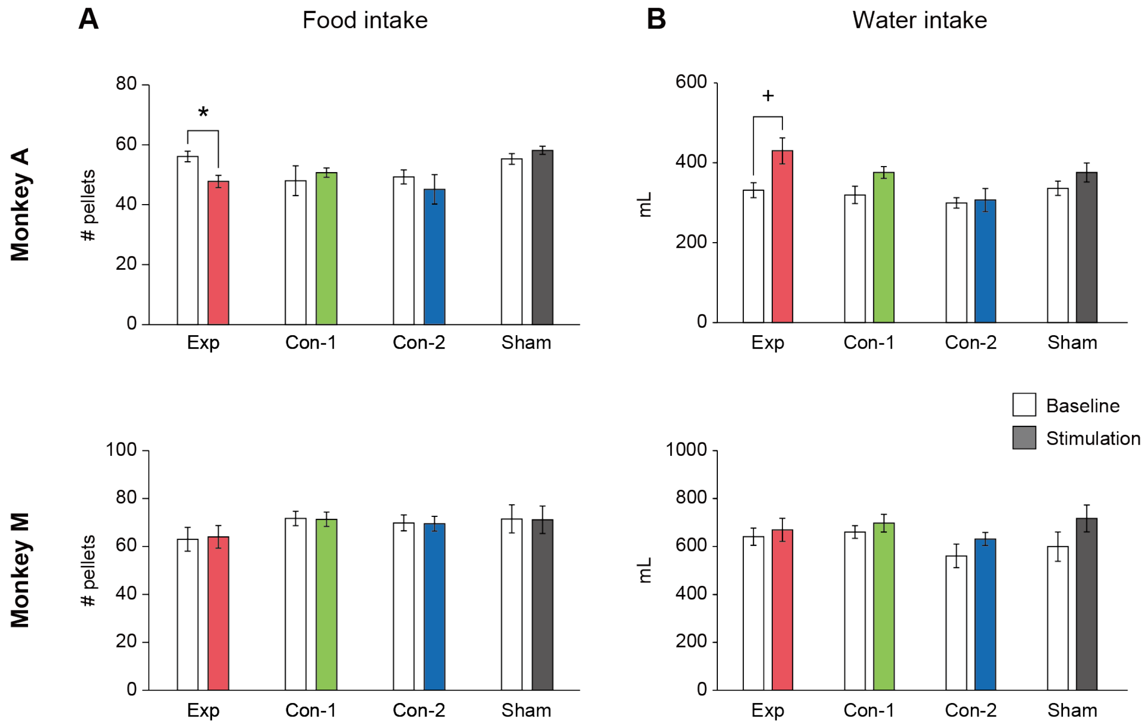


**Fig. S4** Changes in food (A) and water (B) intakes before and after the treatment under each condition. Note that the food intake is shown as the number of pellets (approximately 2.5 g/pellet)) the monkey ate in a day. The data are shown as mean ± SEM. $^{+}$, **P* < 0.1, 0.05, t-test with Bonferroni correction for multiple comparisons.

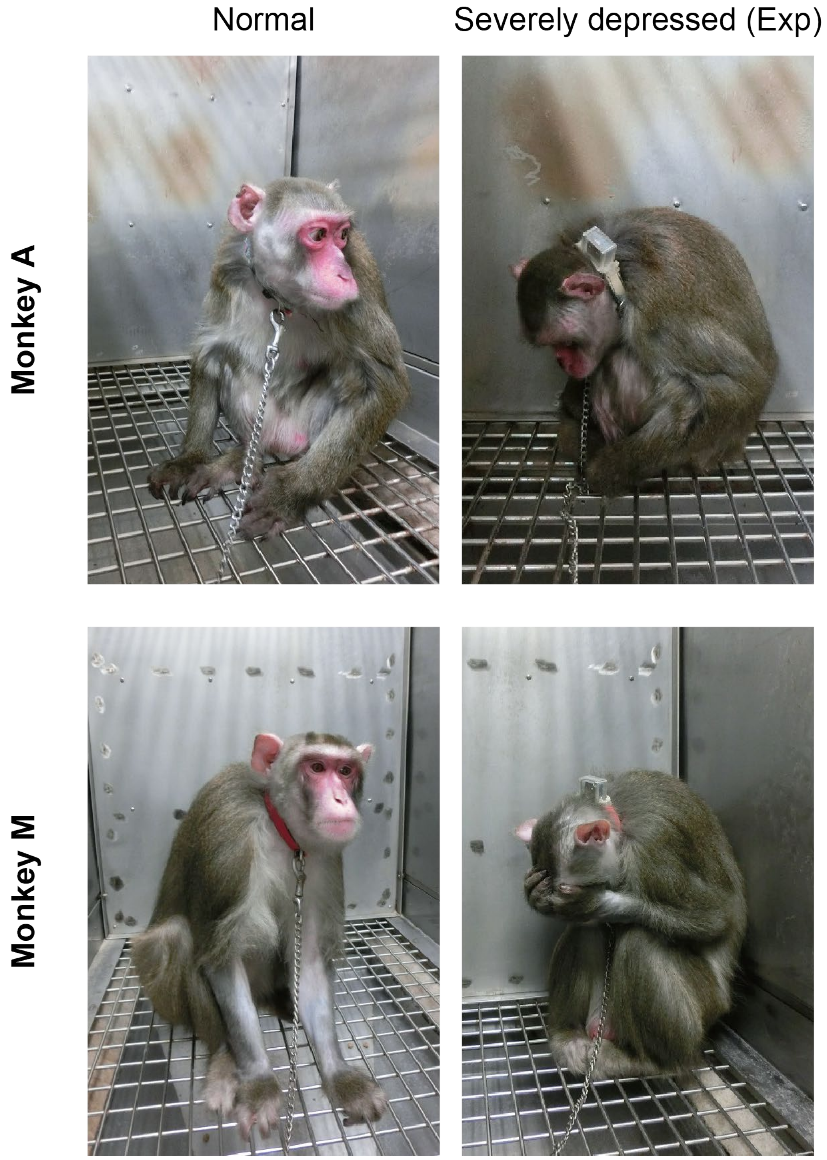


**Fig. S5** Enhanced effect of LF-rTMS on monkeys' behavior. When we conducted multiple sessions (one session/day) of LF-rTMS targeting the vMFC, the monkeys exhibited a severe and prolonged depressive state. They spent much of their time in the back of the cage, leaned against the wall, and looked down with a hunched posture. These symptoms continued for a while, and the monkeys gradually recovered and returned to their normal state about a month later.
